# Genetic characterization of the zoonotic parasite *Ancylostoma caninum* in the central and eastern USA

**DOI:** 10.1101/2022.12.22.521591

**Authors:** Theresa A. Quintana, William L. Johnson, Deb Ritchie, Vicki Smith, Katy A. Martin, Krysta McMahan, Matthew T. Brewer, Jeba R. J. Jesudoss Chelladurai

## Abstract

*Ancylostoma caninum* is the most common nematode parasite of dogs in the United States. This study aimed to describe the molecular epidemiology of *A. caninum* isolates from the central and eastern states of the U.S. using the partial mitochondrial *cox1* gene and to compare them with those reported globally. We isolated eggs from fecal samples of dogs and characterized each isolate based on *cox1* sequences. A total of 60 samples originating from Kansas, Iowa, New York, Florida, and Massachusetts were included. 25 haplotypes were identified in the U.S. dataset with high haplotype diversity (0.904). Sequence data were compared to sequences from other world regions available in GenBank. Global haplotype analysis demonstrated 35 haplotypes with a haplotype diversity of 0.931. Phylogenetic and network analysis provide evidence for the existence of moderate geographical structuring of A. *caninum* haplotypes. Our results provide an updated summary of A. *caninum* haplotypes and data for neutral genetic markers with utility for tracking hookworm populations. Sequences have been deposited in GenBank (ON980650 - ON980674). Further studies of isolates from other regions are essential to understand the genetic diversity of this parasite.

## 1. Introduction

Canine hookworms are important nematodes of dogs worldwide and are causative agents of zoonotic cutaneous larva migrans (CLM) in humans. Of the three canine-adapted hookworm species in the United States, *Ancylostoma caninum, A. braziliense* and *Uncinaria stenocephala, A. caninum* is the most common hookworm in dogs (Bowman *et al.*, 2010). *Ancylostoma ceylanicum*, a zoonotic hookworm endemic in other parts of the world has not been reported from the continental United States.

The prevalence of canine hookworms between 2017 and 2019 in the United States as assessed by fecal floatation and antigen testing in dogs that presented to veterinarians for wellness visits was 4.1%, whereas those that presented for other clinical visits was 4.2% (Sweet *et al.*, 2021). Annual hookworm incidence in dogs across all states increased by 47% between 2015 and 2018 (Drake & Carey, 2019), while some states such as Colorado had an increase of 137% between 2013 to 2017, likely due to dog movement and importation of infected dogs (Drake & Parrish, 2020). Even in large urban areas of the U.S. where 68.8% of dog owners reported providing parasite control medication to their pets, 7.1% of dogs randomly sampled in dog parks were infected with hookworms (Stafford *et al.*, 2020). Additionally, *A. caninum* isolates resistant to multiple anthelmintics are an emerging problem in the U.S. Greyhounds raised in racing kennels are especially prone to be infected by resistant A. *caninum* (Jimenez Castro *et al.*, 2019; Kitchen *et al.*, 2019).

Transmission of hookworms begins when infective A. *caninum* third larval stage (L_3_) penetrates the skin of dogs and humans from dog-feces contaminated environments. The intensity and life cycle of A. *caninum* infections in dogs depend on host age and immunity. In immunocompetent dogs, infection typically begins by ingestion of L_3_ larvae or skin penetration by L_3_ larvae. A proportion of L_3_s can undergo migration within the body to musculature or fat tissue, where they reside as “somatic larvae” in an arrested state of development. Upon endocrine stimulation during the last trimester of pregnancy, these arrested larvae reactivate and can pass vertically to nursing puppies through the mammary glands (Burke & Roberson, 1985). Additionally, “larval leak” may occur, a phenomenon in which reactivated somatic larvae migrate to the lumen of the intestine when existing populations of adult worms in the intestinal niche are eliminated. Thus, infection may be acquired by dogs from many sources. Infections in dogs result in loss of blood due to the hematophagous nature of the pre-adult and adult stages (Stassens *et al.*, 1996). Clinically, blood loss manifests as a spectrum of signs ranging from asymptomatic infections in adult dogs to peracute disease and death in neonates (Hill, 1946; Miller, 1968). Asymptomatic dogs pose the same level of risk as sources of human infections as symptomatic animals (Savilla *et al.*, 2011).

A few records of A. *caninum* reaching patency in humans exists, albeit in a minor fraction of infections in tropical regions (Furtado *et al.*, 2020; George *et al.*, 2016). In a larger proportion of people, larval skin penetration by A. *caninum* produces epidermal migration of larvae which manifests as ephemeral papular, pustular or follicular lesions (Diakou *et al.*, 2019). CLM in humans is often associated with travel and/or close human-animal bonds, especially in areas where anthelmintics are not used in dogs (Chris & Keystone, 2016; Heukelbach *et al.*, 2002). Variability in host and parasite factors have been hypothesized to play a role in the incubation period (Siriez *et al.*, 2010). Parasite factors such as strain variability have been suggested to be related to pathogenicity in the related hookworm *Necator americanus* (Clements & Addis Alene, 2022). Understanding the genetic variability in hookworm populations is crucial to elucidating their epidemiology and has implications for their control (Gasser *et al.*, 2009).

Neutral barcoding genes such as the mitochondrial *cox1* have been deemed to be better genetic markers to understand population structure than nuclear loci such as ITS-1 and ITS-2 (Gasser *et al.*, 2009). There is relatively little data available on the population structure and molecular epidemiology of *A. caninum* globally. In previous studies, 7 haplotypes were identified from 38 adult A. *caninum* from Australia (Hu *et al.*, 2002), 18 haplotypes were identified from 62 U.S. mid-Atlantic samples from Pennsylvania and North Carolina (Moser *et al.*, 2007) and 30 haplotypes were identified from 160 adult worms from Brazil (Miranda *et al.*, 2008). U.S. samples from the other states have not been studied. Additionally, sequences from the previous U.S. study (Moser *et al.*, 2007) are not available in GenBank and there are no publicly available U.S. sequences of the barcoding mitochondrial *cox1* for comparative studies.

The aim of this study was to understand the haplotype distribution, diversity and phylogenetics of A. *caninum* populations isolated from naturally infected dogs in the United States using the partial mitochondrial *cox1* gene and to compare it with those reported globally. We hypothesized that geographical isolation would drive global haplotypic differentiation between A. *caninum* populations in the U.S. and other global regions. We also hypothesized that the genetic structure within the U.S. would be complex due dog movement and importation (Drake & Parrish, 2020).

## 2. Materials and methods

### 2.1. Parasites

Fecal samples of naturally infected, pet dogs were submitted with owner and veterinarian consent to the Kansas State Veterinary Diagnostic Laboratory between December 2020 and April 2022 for diagnostic parasitology testing. *Ancylostoma caninum eggs were* isolated from fecal samples using a double centrifugal fecal flotation technique. Briefly, approximately 5 g of the fecal sample was mixed with water, strained, sedimented with centrifugation and the supernatant was discarded. The sediment was then mixed Sheather’s sugar solution (Sp. gr. 1.275) and a centrifugal floatation was performed with a coverslip covering the tubes. All parasites stages observed on the coverslip were recorded. If *Ancylostoma* spp. eggs were present, the eggs were collected for inclusion in the study.

Three adult A. *caninum* samples were opportunistically obtained from samples submitted for diagnostic identification to Iowa State University College of Veterinary Medicine by veterinarians. All samples were anonymized for genetic studies, except for state of origin and reported dog breed, which were recorded as meta-data. Samples for which the dog breed was not provided were coded as ‘unknown’. Samples used in the final genetic analysis were obtained from: Kansas (51 samples), New York (4 samples), Iowa (3 samples), Florida (1 sample) and Massachusetts (1 sample). All protocols were approved by the Institutional Bio-safety Committee at Kansas State University (IBC-1637-VCS).

### 2.2. DNA extraction, amplification and sequencing of *cox1*

Eggs collected from coverslips (n = 63 samples) were washed using 1X PBS and stored at 4°C until DNA extraction. Adults (n = 3 samples) were stored in 70% ethanol at room temperature until DNA extraction. DNA was extracted from whole adult worms or eggs suspended in a 200μL volume using the Qiagen DNeasy Blood and Tissue kit (Valencia, CA) according to manufacturer’s instructions. Total genomic DNA was eluted in 100 - 200μL of water and stored at −20°C.

An approximately 400 bp fragment of the mitochondrial cytochrome oxidase 1 (*cox1*) gene was amplified using the primers JB3 (5’-TTTTTTGGGCATCCTGAGGTTTAT-3’) and JB4.5 (5’-TAAAGAAAGAACATAATGAAAATG-3’) (Hu *et al.*, 2002). PCR was carried out in a 25μL volume with 2μL of DNA, 1 × PCR buffer, 3mM MgCl_2_, 200 μM of each dNTP, 200nM of each primer and 0.3 units of Taq polymerase (GoTaq Flexi, Promega, Madison, WI). PCR conditions were 94°C for 5 minutes followed by 30 cycles of 94°C for 30s, 55°C for 30 s and 72°C for 30 s, with the final extension of 72°C for 5 min. PCR reactions were analyzed by agarose gel electrophoresis to confirm amplification. Amplicons were enzyme purified (Applied Biosystems, Thermo Fisher Scientific, Vilnius, Lithuania) and sequenced using Sanger sequencing technology (Eurofins Genomics, Louisville, KY). Sequences were analyzed and contigs assembled with GeneStudio ver. 2.2.0.

### 2.3. Sequence analysis

Nucleotide BLAST searches were used to confirm species identity. *A. caninum* nucleotide sequences from (Hu *et al.*, 2002) and others from the same portion of the *cox1* gene were obtained from GenBank for inclusion in this study (Table 1). For phylogenetic analyses, *cox1* sequences of A. *tubaeforme, A. duodenale* and A. *ceylanicum* were also obtained from GenBank for inclusion. *cox1* sequences of A. *braziliense* were not available in the nucleotide database of GenBank for inclusion (Accessed December 19, 2022).

**Table 1.**
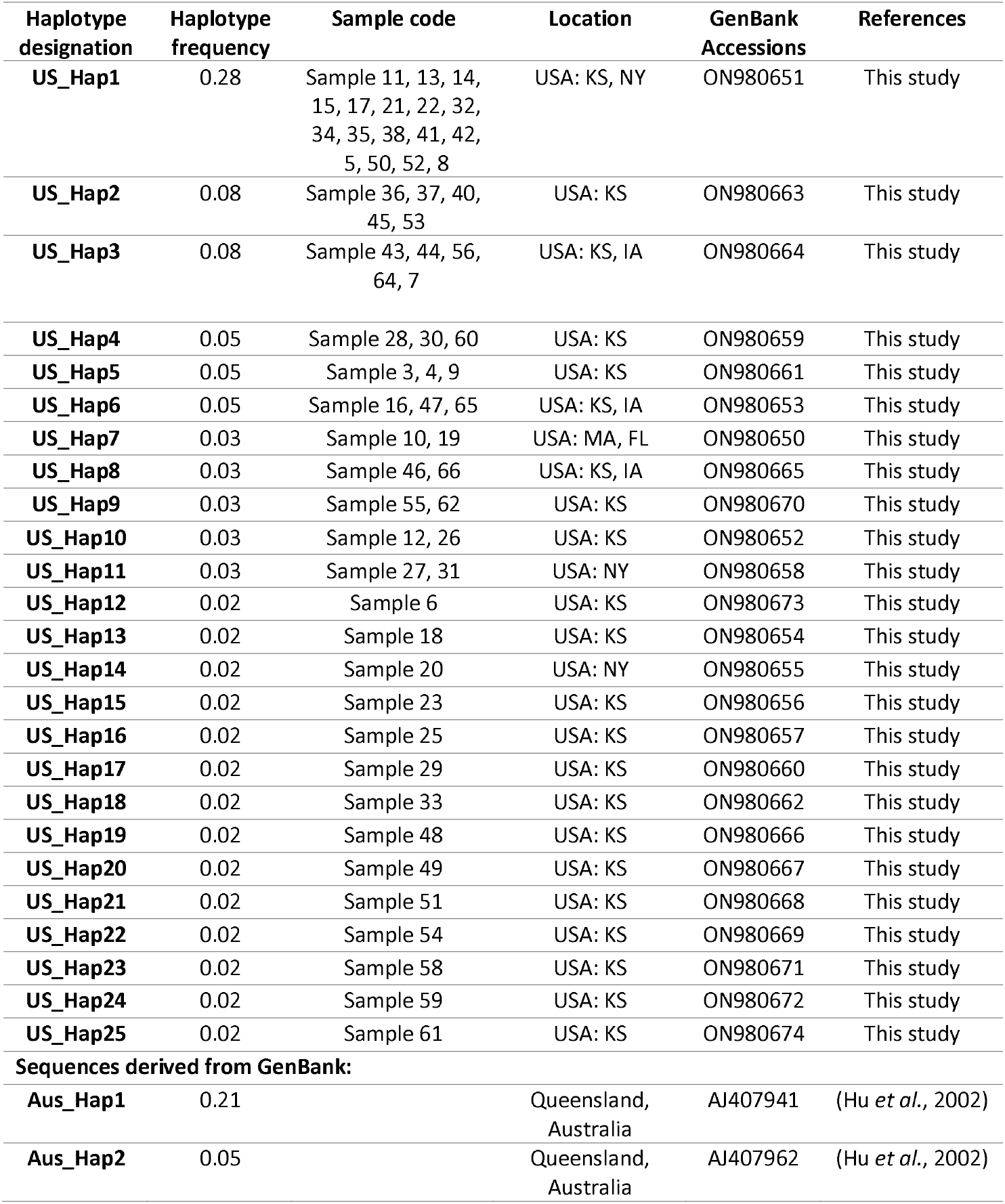
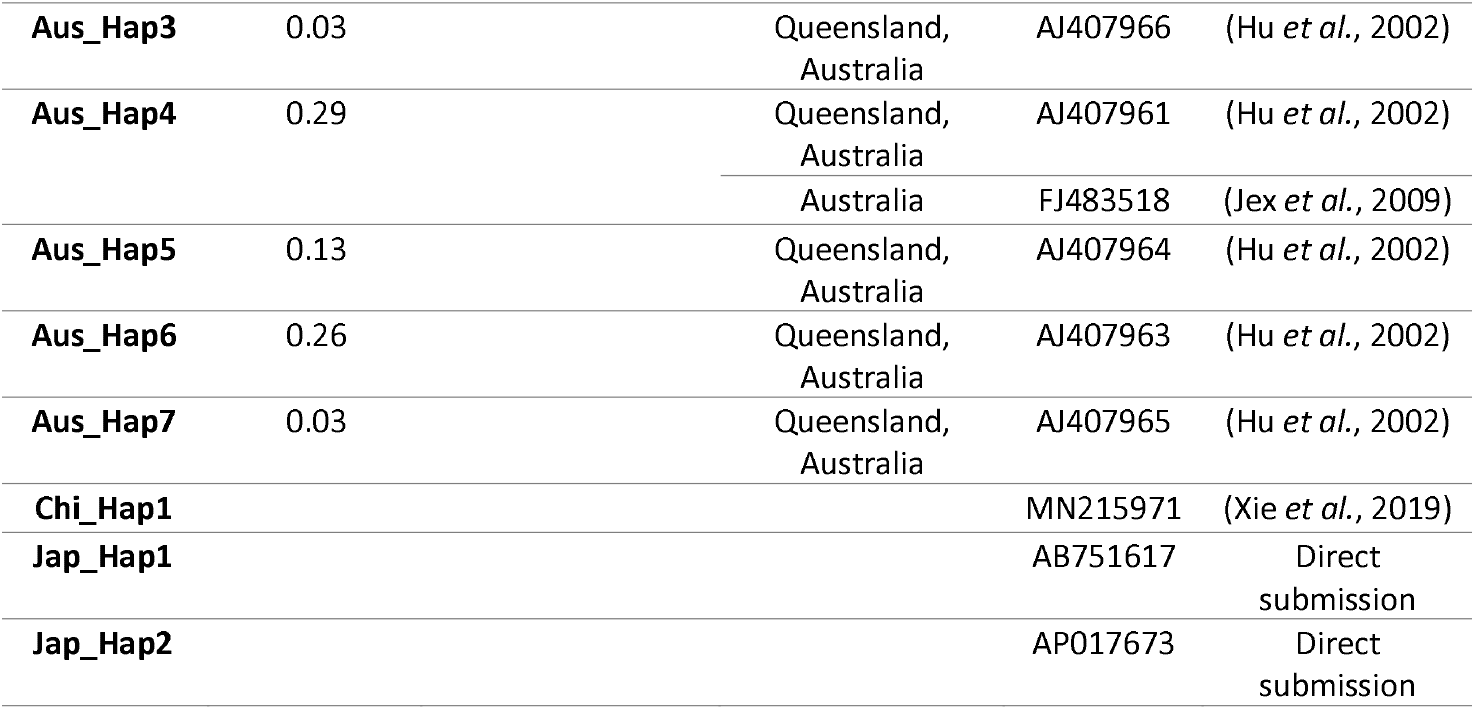
Summary of haplotype analysis of *cox1* sequences obtained from this study and derived from GenBank for comparative analysis.

Multiple sequence alignment was carried out with MAFFT (Katoh & Standley, 2013) and trimmed to 380 bp in MegaX (Kumar *et al.*, 2018) for haplotypic and network analyses, and with BMGE (Criscuolo & Gribaldo, 2010) for phylogenetic analysis. Haplotype diversity, nucleotide diversity and nucleotide difference analyses were performed in DnaSp ver. 6 (Rozas *et al.*, 2017). Geographical distribution of haplotypes were visualized by mapping using datawrapper (Datawrapper, 2021) and Microsoft Excel ver. 16.63. A heatmap of breed distribution against haplotypes in this study was created using plotly (Plotly Technologies, 2015). Median joining networks were generated in PopART (Leigh & Bryant, 2015). Maximum likelihood phylogenetic analysis was performed using PhyML 3.0 (Guindon *et al.*, 2010) after the best substitution model was determined using BIC criteria in SMS (Lefort *et al.*, 2017). The tree was visualized with iTol ver. 6 (Letunic & Bork, 2021).

### 2.4 Data availability

Unique haplotypic sequences generated from this study have been deposited in the GenBank nucleotide database under accession numbers ON980650-ON980674.

## 3. Results

### 3.1 Analysis of *cox1* sequences reveals high haplotypic variability

An approximately 400 bp region of the mitochondrial cytochrome oxidase 1 gene (*cox1*) was amplified from 57 of 63 *A. caninum* positive fecal floats and from 3 of 3 A. *caninum* adult worms. *cox1* sequences obtained from the 60 U.S. samples were found to be distributed across 25 unique haplotypes (Table 1, Figure 1A), with haplotype diversity of 0.904 (variance 0.00085 and standard deviation 0.029) and nucleotide diversity of 0.00886 (standard deviation 0.00086). Haplotype designations were provided to the sequences based on frequency of occurrence. The most frequent haplotype (frequency 0.28) was designated US Hap 1, the next most frequent was designated US Hap 2 etc. As shown in Figure 1A, samples from Kansas (n = 51) represented 22 haplotypes (haplotypic diversity: 0.886 ± 0.04). Sequences from Iowa (n =3) belonged to the same haplotypes as sequences from Kansas (US Hap 3, US Hap 6 and US Hap 8) (haplotypic diversity: 1.0 ± 0.27). Three of the sequences from New York represented two unique haplotypes (US Hap 11 and US Hap 14), while the other belonged to US Hap 1 (haplotypic diversity: 0.83±0.22). Two sequences from Massachusetts and Florida (n = 1 each) belonged to a unique haplotype (US Hap 7).

**Figure 1.**
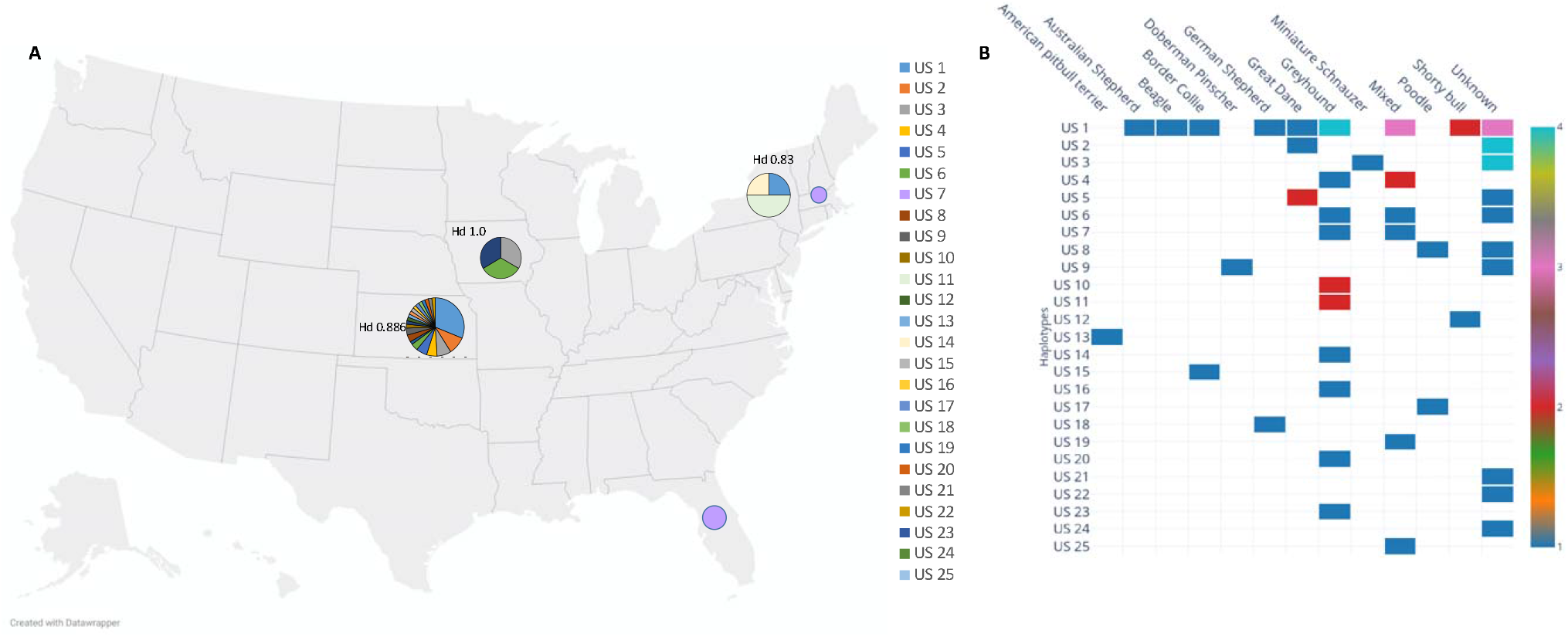
Haplotype distribution of *cox1* sequences from this study. Haplotype diversity is represented as pie charts with analysis performed according to the state of origin of samples. Hd indicates the haplotypic diversity in each state from which >1 sample was analyzed. (B) **Heat map showing haplotype and breed association from this study.**Number of samples belonging to each haplotype derived from *A. caninum* infected dogs of different breeds is represented using a color spectrum (Color blocks blue, red, pink and teal representing 1, 2, 3 and 4 samples respectively).

Global haplotype analysis was performed with 60 sequences from this study, in addition to 11 sequences derived from the nucleotide database of GenBank. Sequences from a study in Brazil (Miranda *et al.*, 2008) were derived from a different portion of the *cox1* gene with no overlap with the sequences in Hu et al. (Hu *et al.*, 2002) and the current study. Sequences from a previous U.S. study (Moser *et al.*, 2007) were not available in GenBank or any other database for inclusion. Haplotype analysis revealed that the 71 sequences in the final dataset represented 35 haplotypes, with a haplotype diversity of 0.931 (variance: 0.00048 and standard deviation: 0.022) and nucleotide diversity of 0.01624 (standard deviation: 0.00303). Of 380 bp analyzed, 59 sites were determined to be polymorphic (variable) with 63 mutations, and 43 sites were determined to be parsimony informative.

The occurrence of haplotypes in different dog breeds included in this study is represented as a heat map (Figure 1B). A diverse set of haplotypes were recorded from greyhounds and unknown breeds, which were both overrepresented in the sample set. The most frequently occurring haplotype US Hap 1 was not restricted to any breed. Unique haplotypes (US Hap 12 – 25) were overrepresented in Greyhounds due to the sampling bias associated with opportunistic sampling.

### 3.2 Haplotype network analysis supports the presence of three clusters

A median joining network of the haplotype data set (n = 35 sequences; each representing 1 haplotype) was constructed. Haplotypic nomenclature, geographical origin, and number of sequences for each haplotype was added from previous studies when available, as summarized in Table 1. Three distinct clusters could be visualized, with all haplotypes from the U.S. and 4 of the 7 haplotypes from Australia forming a cluster (cluster A). Three haplotypes from Australia formed a distinct cluster (cluster B), which had an average nucleotide difference of 27.6 from cluster A. Fst between haplotypes in Cluster A and B was 0.80. Fst between all U.S. haplotypes and all Australian haplotypes (in both clusters A and B) was 0.249. A single GenBank sequence from China was distinct (designated Cluster C) with an average nucleotide difference of 16.9 from cluster A. Clusters B and C had an average difference of 32.7 nucleotides. Fst between cluster C and other clusters could not be calculated because cluster C comprised of a single sequence. Additionally, there is some evidence for a sub cluster within cluster A formed by the haplotypes US Hap12, US Hap 18, Aus Hap 7, and Jap Hap1. Fst and average number of nucleotide differences between the subcluster population and all the other haplotypes of cluster A is 0.335 and 8.8 respectively. Taken together, this data suggests there is moderate genetic structuring of global A. *caninum* populations.

### 3.3 Phylogenetic analysis supports the existence of genetic clusters

A maximum likelihood tree of unique haplotypes was constructed using the GTR model for nucleotide substitutions with discrete gamma model parameters (gamma classes: 4, shape parameter: 0.057) and bootstrap branch supports (Figure 3). *Cooperia oncophora cox1* sequence (GenBank Accession: GQ888713) was used as the outgroup. All sequences of A. *caninum* formed a monophyletic clade with high statistical support (91%). Sequences from the U.S. in the study were in Cluster A, which appeared monophyletic but with low statistical support (32%). Three haplotypic sequences from Australia formed a distinct cluster (Cluster B) with high statistical support (96%), but this cluster was monophyletic with the U.S. isolates with 64% statistical support. The single *cox1* sequence from China formed a distinct cluster (Cluster C) but had low statistical support (<50%). The identity of A. *caninum* sequences from this study as being distinct from the sequences of other *Ancylostoma* spp. was highly supported by the monophyly of A. *tubaeforme* (100%), A. *duodenale* (84%) and A. *ceylanicum* (99%).

**Figure 2.**
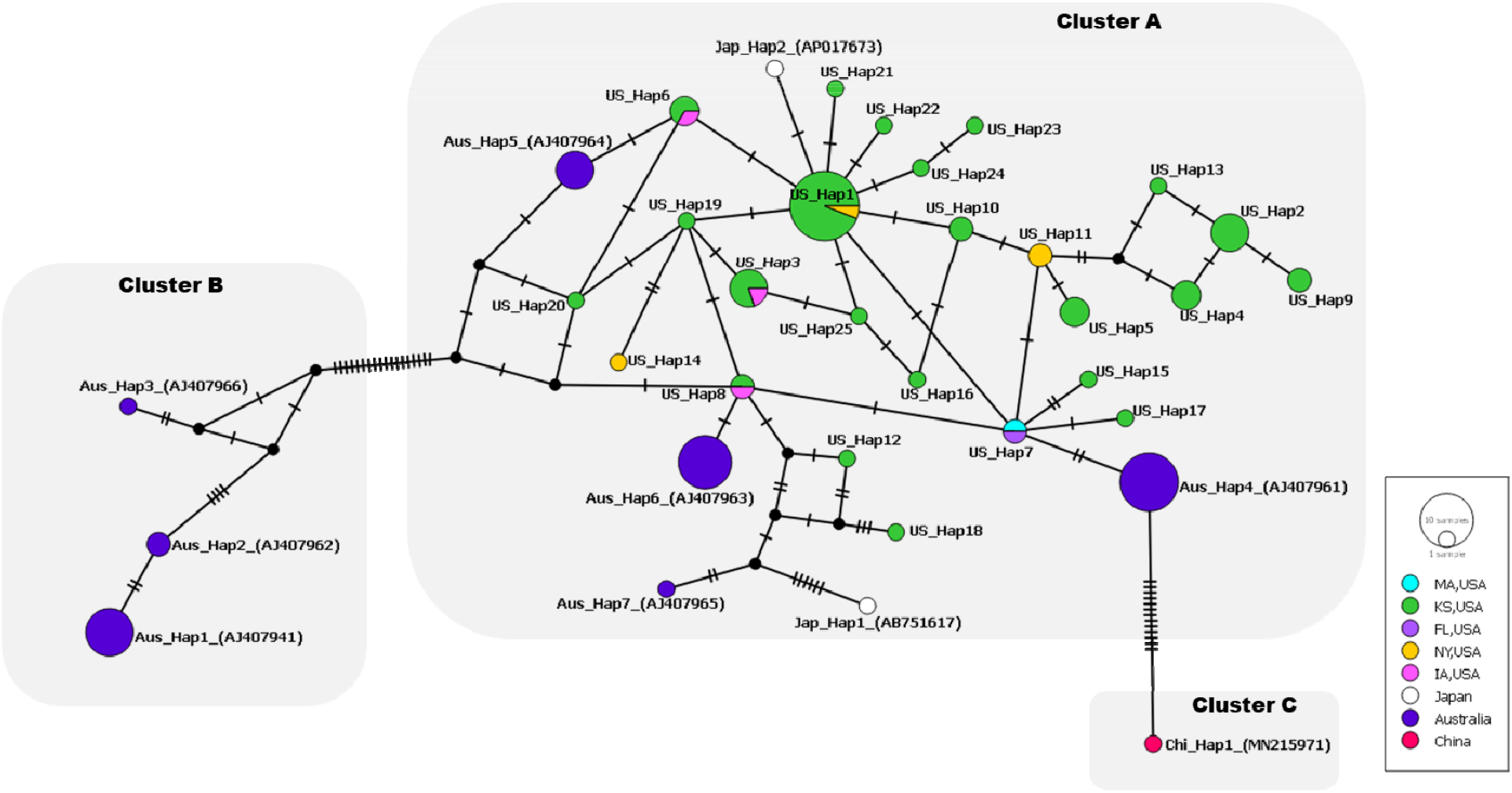
Median joining network of *cox 1* haplotypes from this study (designated US Hap 1 – 25) and GenBank sequences (denoted by country, haplotype designation, and accession numbers). Haplotype circles are colored to represent unique geographical sources of the sequence and are scaled to represent the number of sequences belonging to each haplotype (this study and (Hu et al., 2002)). Nucleotide differences are denoted by hatch marks across the connecting lines with each mark representing a single nucleotide difference. Unlabeled dark circles represent inferred, unsampled nodes. Three clusters observable in the network are represented by grey boxes.

**Figure 3.**
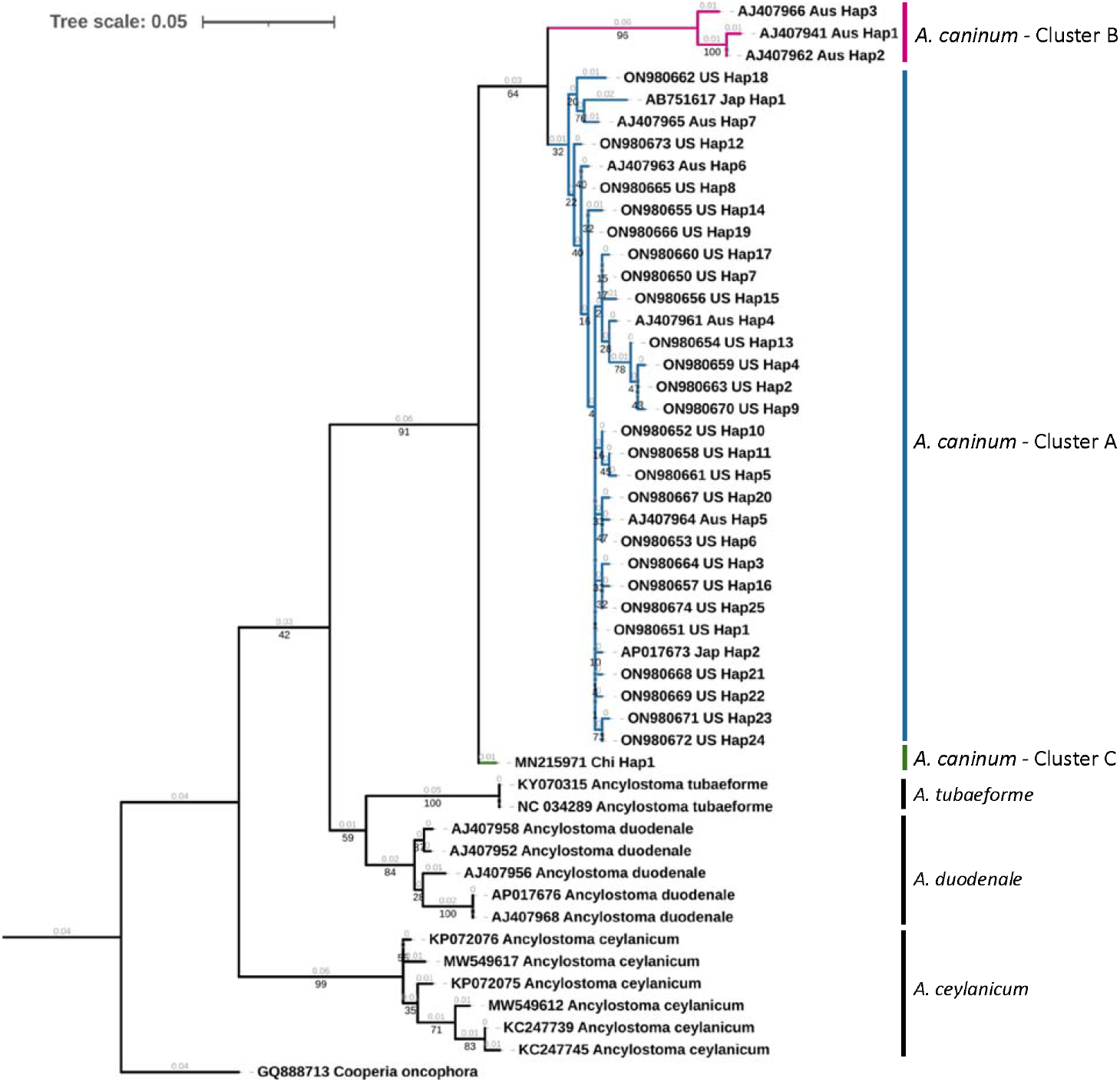
Phylogenetic tree of *Ancylostoma* spp. A maximum likelihood tree of haplotypes generated from *A. caninum* partial *cox 1* sequences in this study and *Ancylostoma* spp. sequences derived from GenBank. Accession numbers, country code and haplotypic designation are given for each haplotype. *Cooperia oncophora cox1*, trimmed from the complete mitochondrial genome record (Accession number: GQ888713), was used as the outgroup. Cluster designations based on the haplotype network (Figure 2) are provided. Bootstrap values are represented in black under branches. Computed branch lengths are represented in grey above the branches.

## 4. Discussion

*Ancylostoma caninum* is one of the most common nematode parasites of dogs in the United States. As a causative agent of human CLM, its prevalence in dogs, incidence, genetic variation, and viability in the environment, among other factors are important from a One Health perspective. Incidence of CLM can be reduced by proper anthelmintic administration to dogs and by removing dog feces immediately after defecation after donning appropriate PPE and instituting hygiene practices. Understanding the molecular epidemiology in dogs is an crucial starting point for the control of the infection in humans since prevalence and differences in pathogenicity in CLM at the population level is unknown (Heukelbach *et al.*, 2002). Yet, there is an important need for this information due to the reported increase in the incidence of infection in dogs, which increases the risk of human infection. We hypothesized that the geographical isolation of A. *caninum* populations would lead to the emergence of novel haplotypes in the U.S. and that genetic structure within the U.S. would be complex due to dog movements. To test these hypotheses, we examined *A. caninum* in dog isolates from the U.S. and characterized prevalent mitochondrial cox1 haplotypes.

The mitochondrial *cox1* has been used as a neutral barcoding marker in multiple parasite studies (Jesudoss Chelladurai *et al.*, 2017; Small *et al.*, 2014). Ideally, molecular epidemiology studies must use published protocols if any exist (such as using the same primer sets) and must deposit sequence data generated in an easily accessible public database. This approach has resulted in a plethora of phylogeographic epidemiological data being generated for important zoonotic parasites - *Ascaris* spp. and *Echinococcus* spp. (Cavallero *et al.*, 2013; Spotin *et al.*, 2018). In the case of *A. caninum*, however, genetic data is scant due to unavailability of data from previous studies and is complicated by the use of different primer sets by different research groups. The complete *cox1* gene is 1577 bp long in the *A. caninum* reference mitochondrial genome (GenBank Accession: NC_012309). The primer set used by Hu et al. (Hu *et al.*, 2002) flank nucleotides 754 to 1146 of the *cox1*, while the primer set used by Miranda et al. (Miranda *et al.*, 2008) flanks nucleotides 162 to 628 and the primer set used by Mulinge et al. (Mulinge *et al.*, 2021) flanks nucleotides 368 to 830. In this study, the primers used by (Hu *et al.*, 2002) and (Moser *et al.*, 2007) were used for PCR amplification.

All haplotypes identified in this study were unique to the United States. Our analysis revealed 35 unique *A. caninum* haplotypes currently reported globally and 25 unique haplotypes within the U.S. Until now, there has been a paucity of publicly available molecular data from *A. caninum* populations in the United States. Unique haplotypes were also found in a previous U.S. study (Moser *et al.*, 2007), but the absence of publicly available data from that study makes direct comparisons impossible. Overall haplotypic diversity among U.S. samples in this study (0.931; 60 samples) was higher than diversity at state level – 0.886 in Kansas (51 samples) and 0.833 in New York (4 samples; Figure 1A). The higher diversity (1.0) reported from Iowa is attributable to the small random sample obtained for inclusion in the study (3 samples). State level haplotype diversities from this study were comparable to A. *caninum* diversity previously reported from Australia (0.80; 38 samples) (Hu *et al.*, 2002), from North Carolina, USA (0.828; 54 samples) (Moser *et al.*, 2007) and Brazil (0.88; 164 samples) (Miranda *et al.*, 2008). These diversities are significantly higher than the A. *caninum* diversity reported from Massachusetts, USA (0.25; 8 samples) (Moser *et al.*, 2007) and diversity of other soil-transmitted zoonotic nematodes like *Ascaris* from U.S. populations (0.596; 100 samples) (Jesudoss Chelladurai *et al.*, 2017). Additionally, high numbers of A. *caninum* haplotypes were recorded in greyhounds, a breed overrepresented in reports of anthelmintic resistance (Figure 1B).

Median joining haplotype network analysis demonstrated a complex pattern (Figure 2). A. *caninum* occurs as three clusters, in agreement with (Moser *et al.*, 2007). U.S. haplotypes were found only in cluster A, while Australian haplotypes were found in clusters A and B. Cluster B currently only has haplotypes from Australia and cluster C has only 1 haplotype from China. Combined with the Fst values obtained, it can be concluded that there is moderate population structuring in A. *caninum.* The maximum likelihood tree provided high statistical support for haplotypes in cluster B (Figure 3). Due to the moderately high statistical support for the supercluster containing clusters A and B (64%), it is evident that the sequence in cluster C is distinct from clusters A and B.

Haplotype differentiation and spread can be attributable to many factors such as hybridization, introgression and retention of ancestral polymorphisms (Detwiler & Criscione, 2010). Dog movement and dog importation also play a role in the spread of zoonotic parasites (von Rentzell *et al.*, 2022; Wright *et al.*, 2020). Rates of mutation that led to the formation of novel haplotypes is still relatively unknown in zoonotic helminths (Jesudoss Chelladurai *et al.*, 2017).

Some limitations of this study include a geographical sampling bias associated with the selection of A. *caninum* sent to a state veterinary diagnostic lab. Since samples were opportunistically collected, clinical history was not considered as a factor in the analysis and often not provided by submitters. The use of Sanger sequencing allowed the capture of only the predominant cox1 haplotype in each sample. Additionally, the use of cox1 primers from Hu et al. (Hu *et al.*, 2002) disallowed us from making comparisons with the dataset from (Miranda *et al.*, 2008). Additionally, the lack of A. *caninum* gene sequences from across the world in GenBank and/or other public nucleotide databases led to a low global representation of A. *caninum* populations in this study. These limitations can be overcome in future studies by prospective selection of dogs with adequate clinical history for inclusion in studies and by the use of deep amplicon sequencing to detect intra-sample variations in mitochondrial sequences. Molecular tools can be useful in increasing our understanding of A. *caninum* and therefore CLM epidemiology. The usefulness of cox1 sequences amplified from eggs in describing infection epidemiology has been previously demonstrated with human hookworms (Monteiro *et al.*, 2019). In the present study we provide molecular characterization of A. *caninum* from dogs in the United States. Sequences from this study are available in GenBank for future comparative analyses. Our results demonstrated that dogs harbor haplotypes of A. *caninum* unique to the U.S. Future studies of A. *caninum* isolates from endemic regions across the world are essential to understand the genetic diversity of this parasite.

## Acknowledgements

The authors would like to acknowledge the help provided by Chance Kopsa, Dr. Brian Herrin, Dr. Kamilyah Miller, and Dr. Cameron Sutherland for help with holding and storing of hookworm positive samples after diagnostic work was completed at the KSVDL.

## Meeting where data was presented

A portion of the data was presented at the 66th Annual American Association of Veterinary Parasitologists (AAVP Hybrid Meeting – Lexington, KY and virtual). June 2021.

## CRediT authorship

Conceptualization: J.R.J.; Data curation: T.A.Q. and J.R.J.; Formal analysis: T.A.Q. and J.R.J.; Funding acquisition: J.R.J.; Investigation: T.A.Q., W.L.J., D.R., V.S., K.A.M., K.M., M.T.B. and J.R.J.; Project administration: J.R.J.; Resources: M.T.B. and J.R.J.; Supervision: J.R.J.; Visualization: T.A.Q. and J.R.J.; Writing – original draft: J.R.J.; Writing - review & editing: T.A.Q., W.L.J., D.R., V.S., K.A.M., K.M., M.T.B. and J.R.J.5.

## References

Bowman, D.D., Montgomery, S.P., Zajac, A.M., Eberhard, M.L. & Kazacos, K.R. (2010) Hookworms of dogs and cats as agents of cutaneous larva migrans. Trends Parasitol, 26, 162–167. doi: 10.1016/j.pt.2010.01.005.

Burke, T.M. & Roberson, E.L. (1985) Prenatal and lactational transmission of Toxocara canis and Ancylostoma caninum: experimental infection of the bitch at midpregnancy and at parturition. Int J Parasitol, 15, 485–490. doi: 10.1016/0020-7519(85)90041-4.

Cavallero, S., Snabel, V., Pacella, F., Perrone, V. & D’Amelio, S. (2013) Phylogeographical studies of Ascaris spp. based on ribosomal and mitochondrial DNA sequences. PLoS Negl Trop Dis, 7, e2170. doi: 10.1371/journal.pntd.0002170.

Chris, R.B. & Keystone, J.S. (2016) Prolonged incubation period of Hookworm-related cutaneous larva migrans. J Travel Med, 23, tav021. doi: 10.1093/jtm/tav021.

Clements, A.C.A. & Addis Alene, K. (2022) Global distribution of human hookworm species and differences in their morbidity effects: a systematic review. Lancet Microbe, 3, e72–e79. doi: 10.1016/S2666-5247(21)00181-6.

Criscuolo, A. & Gribaldo, S. (2010) BMGE (Block Mapping and Gathering with Entropy): a new software for selection of phylogenetic informative regions from multiple sequence alignments. BMC Evol Biol, 10, 210. doi: 10.1186/1471-2148-10-210.

Datawrapper (2021) Datawrapper tool. https://www.datawrapper.de. Secondary Datawrapper tool, https://www.datawrapper.de. www.datawrapper.de/ 2021.

Detwiler, J.T. & Criscione, C.D. (2010) An infectious topic in reticulate evolution: introgression and hybridization in animal parasites. Genes (Basel), 1, 102–123. doi: 10.3390/genes1010102.

Diakou, A., Di Cesare, A., Morelli, S., Colombo, M., Halos, L., Simonato, G., Tamvakis, A., Beugnet, F., Paoletti, B. & Traversa, D. (2019) Endoparasites and vector-borne pathogens in dogs from Greek islands: Pathogen distribution and zoonotic implications. PLoS Negl Trop Dis, 13, e0007003. doi: 10.1371/journal.pntd.0007003.

Drake, J. & Carey, T. (2019) Seasonality and changing prevalence of common canine gastrointestinal nematodes in the USA. Parasit Vectors, 12, 430. doi: 10.1186/s13071-019-3701-7.

Drake, J. & Parrish, R. (2020) Dog importation and changes in canine intestinal nematode prevalence in Colorado, USA, 2013-2017. Parasit Vectors, 13, 404. doi: 10.1186/s13071-020-04283-z.

Furtado, L.F.V., Dias, L.T.O., Rodrigues, T.O., Silva, V.J.D., Oliveira, V.N.G.M. & Rabelo, É. (2020) Egg genotyping reveals the possibility of patent Ancylostoma caninum infection in human intestine. Sci Rep, 10, 3006. doi: 10.1038/s41598-020-59874-8.

Gasser, R.B., Cantacessi, C. & Campbell, B.E. (2009) Improved molecular diagnostic tools for human hookworms. Expert Rev Mol Diagn, 9, 17–21. doi: 10.1586/14737159.9.1.17.

George, S., Levecke, B., Kattula, D., Velusamy, V., Roy, S., Geldhof, P., Sarkar, R. & Kang, G. (2016) Molecular Identification of Hookworm Isolates in Humans, Dogs and Soil in a Tribal Area in Tamil Nadu, India. PLoS Negl Trop Dis, 10, e0004891. doi: 10.1371/journal.pntd.0004891.

Guindon, S., Dufayard, J.F., Lefort, V., Anisimova, M., Hordijk, W. & Gascuel, O. (2010) New algorithms and methods to estimate maximum-likelihood phylogenies: assessing the performance of PhyML 3.0. Syst Biol, 59, 307–321. doi: 10.1093/sysbio/syq010.

Heukelbach, J., Mencke, N. & Feldmeier, H. (2002) Editorial: Cutaneous larva migrans and tungiasis: the challenge to control zoonotic ectoparasitoses associated with poverty. Trop Med Int Health, 7, 907–910. doi: 10.1046/j.1365-3156.2002.00961.x.

Hill, H.C. (1946) Observations on Ancylostoma and Toxocara infection in experimental and stock dogs. J Parasitol, 32, 210.

Hu, M., Chilton, N.B., Zhu, X. & Gasser, R.B. (2002) Single-strand conformation polymorphism-based analysis of mitochondrial cytochrome c oxidase subunit 1 reveals significant substructuring in hookworm populations. Electrophoresis, 23, 27–34. doi: 10.1002/1522-2683(200201)23:1<27::AID-ELPS27>3.0.CO;2-7.

Jesudoss Chelladurai, J., Murphy, K., Snobl, T., Bader, C., West, C., Thompson, K. & Brewer, M.T. (2017) Molecular Epidemiology of Ascaris Infection Among Pigs in Iowa. J Infect Dis, 215, 131–138. doi: 10.1093/infdis/jiw507.

Jex, A.R., Waeschenbach, A., Hu, M., van Wyk, J.A., Beveridge, I., Littlewood, D.T. & Gasser, R.B. (2009) The mitochondrial genomes of Ancylostoma caninum and Bunostomum phlebotomum--two hookworms of animal health and zoonotic importance. BMC Genomics, 10, 79. doi: 10.1186/1471-2164-10-79.

Jimenez Castro, P.D., Howell, S.B., Schaefer, J.J., Avramenko, R.W., Gilleard, J.S. & Kaplan, R.M. (2019) Multiple drug resistance in the canine hookworm Ancylostoma caninum: an emerging threat? Parasit Vectors, 12, 576. doi: 10.1186/s13071-019-3828-6.

Katoh, K. & Standley, D.M. (2013) MAFFT multiple sequence alignment software version 7: improvements in performance and usability. Mol Biol Evol, 30, 772–780. doi: 10.1093/molbev/mst010.

Kitchen, S., Ratnappan, R., Han, S., Leasure, C., Grill, E., Iqbal, Z., Granger, O., O’Halloran, D.M. & Hawdon, J.M. (2019) Isolation and characterization of a naturally occurring multidrug-resistant strain of the canine hookworm, Ancylostoma caninum. Int J Parasitol, 49, 397–406. doi: 10.1016/j.ijpara.2018.12.004.

Kumar, S., Stecher, G., Li, M., Knyaz, C. & Tamura, K. (2018) MEGA X: Molecular Evolutionary Genetics Analysis across Computing Platforms. Mol Biol Evol, 35, 1547–1549. doi: 10.1093/molbev/msy096.

Lefort, V., Longueville, J.E. & Gascuel, O. (2017) SMS: Smart Model Selection in PhyML. Mol Biol Evol, 34, 2422–2424. doi: 10.1093/molbev/msx149.

Leigh, J.W. & Bryant, D. (2015) POPART: full-feature software for haplotype network construction. Methods in Ecology and Evolution, 6, 1110–1116.

Letunic, I. & Bork, P. (2021) Interactive Tree Of Life (iTOL) v5: an online tool for phylogenetic tree display and annotation. Nucleic Acids Res, 49, W293–W296. doi: 10.1093/nar/gkab301.

Miller, T.A. (1968) Pathogenesis and immunity in hookworm infection. Trans R Soc Trop Med Hyg, 62, 473–489. doi: 10.1016/0035-9203(68)90130-2.

Miranda, R.R., Tennessen, J.A., Blouin, M.S. & Rabelo, E.M. (2008) Mitochondrial DNA variation of the dog hookworm Ancylostoma caninum in Brazilian populations. Vet Parasitol, 151, 61–67. doi: 10.1016/j.vetpar.2007.09.027.

Monteiro, K.J.L., Jaeger, L.H., Nunes, B.C., Calegar, D.A., Reis, E.R.C.D., Bacelar, P.A.A., Santos, J.P.D., Bóia, M.N. & Carvalho-Costa, F.A. (2019) Mitochondrial DNA reveals species composition and phylogenetic relationships of hookworms in northeastern Brazil. Infect Genet Evol, 68, 105–112. doi: 10.1016/j.meegid.2018.11.018.

Moser, J.M., Carbone, I., Arasu, P. & Gibson, G. (2007) Impact of population structure on genetic diversity of a potential vaccine target in the canine hookworm (Ancylostoma caninum). J Parasitol, 93, 796–805. doi: 10.1645/GE-1096R.1.

Mulinge, E., Zeyhle, E., Mpario, J., Mugo, M., Nungari, L., Ngugi, B., Sankale, B., Gathura, P., Magambo, J. & Kachani, M. (2021) A survey of intestinal helminths in domestic dogs in a human–animal–environmental interface: the Oloisukut Conservancy, Narok County, Kenya. Journal of Helminthology, 95.

Plotly Technologies, I. (2015) Collaborative data science Secondary Collaborative data science. https://plot.lv 2022.

Rozas, J., Ferrer-Mata, A., Carlos Sanchez-DelBarrio, J., Guirao-Rico, S., Librado, P., Ramos-Onsins, S.E. & Sanchez-Gracia, A. (2017) DnaSP 6: DNA Sequence Polymorphism Analysis of Large Data Sets. Molecular Biology and Evolution, 34, 3299–3302. doi: 10.1093/molbev/msx248.

Savilla, T.M., Joy, J.E., May, J.D. & Somerville, C.C. (2011) Prevalence of dog intestinal nematode parasites in south central West Virginia, USA. Vet Parasitol, 178, 115–120. doi: 10.1016/j.vetpar.2010.12.034.

Siriez, J.Y., Angoulvant, F., Buffet, P., Cleophax, C. & Bourrat, E. (2010) Individual variability of the cutaneous larva migrans (CLM) incubation period. Pediatr Dermatol, 27, 211–212. doi: 10.1111/j.1525-1470.2010.01107.x.

Small, S.T., Tisch, D.J. & Zimmerman, P.A. (2014) Molecular epidemiology, phylogeny and evolution of the filarial nematode Wuchereria bancrofti. Infect Genet Evol, 28, 33–43. doi: 10.1016/j.meegid.2014.08.018.

Spotin, A., Boufana, B., Ahmadpour, E., Casulli, A., Mahami-Oskouei, M., Rouhani, S., Javadi-Mamaghani, A., Shahrivar, F. & Khoshakhlagh, P. (2018) Assessment of the global pattern of genetic diversity in Echinococcus multilocularis inferred by mitochondrial DNA sequences. Vet Parasitol, 262, 30–41. doi: 10.1016/j.vetpar.2018.09.013.

Stafford, K., Kollasch, T.M., Duncan, K.T., Horr, S., Goddu, T., Heinz-Loomer, C., Rumschlag, A. J., Ryan, W.G., Sweet, S. & Little, S.E. (2020) Detection of gastrointestinal parasitism at recreational canine sites in the USA: the DOGPARCS study. Parasit Vectors, 13, 275. doi: 10.1186/s13071-020-04147-6.

Stassens, P., Bergum, P.W., Gansemans, Y., Jespers, L., Laroche, Y., Huang, S., Maki, S., Messens, J., Lauwereys, M., Cappello, M., Hotez, P.J., Lasters, I. & Vlasuk, G.P. (1996) Anticoagulant repertoire of the hookworm Ancylostoma caninum. Proc Natl Acad Sci USA, 93, 2149–2154. doi: 10.1073/pnas.93.5.2149.

Sweet, S., Hegarty, E., McCrann, D.J., Coyne, M., Kincaid, D. & Szlosek, D. (2021) A 3-year retrospective analysis of canine intestinal parasites: fecal testing positivity by age, U.S. geographical region and reason for veterinary visit. Parasit Vectors, 14, 173. doi: 10.1186/s13071-021-04678-6.

von Rentzell, K.A., van Haaften, K., Morris, A. & Protopopova, A. (2022) Investigation into owner-reported differences between dogs born in versus imported into Canada. PLoS One, 17, e0268885. doi: 10.1371/journal.pone.0268885.

Wright, I., Jongejan, F., Marcondes, M., Peregrine, A., Baneth, G., Bourdeau, P., Bowman, D.D., Breitschwerdt, E.B., Capelli, G., Cardoso, L., Dantas-Torres, F., Day, M.J., Dobler, G., Ferrer, L., Gradoni, L., Irwin, P., Kempf, V.A.J., Kohn, B., Krämer, F., Lappin, M., Madder, M., Maggi, R.G., Maia, C., Miró, G., Naucke, T., Oliva, G., Otranto, D., Pennisi, M.G., Penzhorn, B.L., Pfeffer, M., Roura, X., Sainz, A., Shin, S., Solano-Gallego, L., Straubinger, R.K., Tasker, S., Traub, R. & Little, S. (2020) Parasites and vector-borne diseases disseminated by rehomed dogs. Parasit Vectors, 13, 546. doi: 10.1186/s13071-020-04407-5.

Xie, Y., Xu, Z., Zheng, Y., Li, Y., Liu, Y., Wang, L., Zhou, X., Zuo, Z., Gu, X. & Yang, G. (2019) The mitochondrial genome of the dog hookworm. Mitochondrial DNA B Resour, 4, 3002–3004. doi: 10.1080/23802359.2019.1666048.

